# Two Piwis with Ago-like functions silence somatic genes at the chromatin level

**DOI:** 10.1101/2020.08.24.263970

**Authors:** Franziska Drews, Sivarajan Karunanithi, Ulrike Götz, Simone Marker, Raphael deWijn, Marcello Pirritano, Angela M. Rodrigues-Viana, Martin Jung, Gilles Gasparoni, Marcel H. Schulz, Martin Simon

## Abstract

Most sRNA biogenesis mechanisms involve either RNAseIII cleavage or ping-pong amplification by different Piwi proteins harboring slicer activity. Here, we follow the question why the mechanism of transgene-induced silencing in the ciliate *Paramecium* needs both Dicer activity and two Ptiwi proteins. This pathway involves primary siRNAs produced from non-translatable transgenes and secondary siRNAs from endogenous remote loci. Our data does not indicate any signatures from ping-pong amplification but Dicer cleavage of long dsRNA. We show that Ptiwi13 and 14 have different preferences for primary and secondary siRNAs but do not load them mutually exclusive. Both Piwis enrich for antisense RNAs and Ptiwi14 loaded siRNAs show a 5′-U signature. Both Ptiwis show in addition a general preference for Uridine-rich sRNAs along the entire sRNA length. Our data indicates both Ptiwis and 2’-O-methylation to contribute to strand selection of Dicer cleaved siRNAs. This unexpected function of two distinct vegetative Piwis extends the increasing knowledge of the diversity of Piwi functions in diverse silencing pathways. As both Ptiwis show differential subcellular localisation, Ptiwi13 in the cytoplasm and Ptiwi14 in the vegetative macronucleus, we conclude that cytosolic and nuclear silencing factors are necessary for efficient chromatin silencing.

## 1 Introduction

RNA silencing is a rather general term for a broad variety of mechanisms regulating gene expression by short RNA molecules. These can either target already transcribed mRNAs post transcriptionally (PTGS) or they can interfere in transcription via co-transcriptional targeting of nascent transcripts (CTGS) thus recruiting chromatin modifying complexes [9, 19]. An important component of any RNAi (RNAinterference) mechanism are Argonaute proteins (Ago) which specifically load sRNAs and mediate further action of those, e.g. by screening cellular RNAs for complementary sequences. Agos themselves can be phylogenetically dissected into two clades: Agos and Piwis (P-element induced wimpy testes), the latter being discovered in *Drosophila* germline stem cells. Agos form the RISC (RNA induced silencing complex) with miRNAs and siRNAs both being ubiquitously expressed, whereas Piwis and their associated piRNAs are expressed in germline cells, only [55].

siRNAs are a distinct class of regulatory RNAs, produced by the dsRNA (double stranded RNA)-specific ribonuclease Dicer: these have been shown across kingdoms to act either in PTGS and CTGS of protein coding genes and structural elements such as centromeres through the life cycle. In many systems, transitivity has been shown by secondary siRNAs involving activity of RNA-dependent-RNA polymerases (RDR).

In contrast to Dicer cleaved siRNAs, piRNAs were described mainly to silence transposable elements during gametogenesis. However, increasing data on different piRNA mechanism reveal an unexpected diversity of those either concerning their targets, their temporal/spatial occurrence in germline and soma and their piRNA biogenesis mechanisms. A similar but far not identical mechanism was described for mouse and *Drosophila*: single stranded precursor 5′-U RNAs loaded into Piwi proteins and this is followed by subsequent 3′-trimming of the piRNA end. This initiation is then followed by Dicer independent amplification of piRNAs by the ping-pong mechanism involving the reciprocal cleavage of complementary ssRNA thus generating a internal single nucleotide A-preference (reviewed in [49]). In all systems, 3′-trimming of piRNAs end is followed by 2′-O-methylation of the mature piRNAs ultimate 3′-nucleotide, this modification is therefore a prominent biochemical mark of piRNAs [54]. The striking difference in piRNA biogenesis by Piwis in contrast to the siRNAs and miRNAs is the difference between Agos and Piwis. Agos load duplexes of dicer cuts and select for guide and passenger strand before screening for targets. In contrast, Piwis are specific for ssRNA: they select those with strong preference for the 5′-U and mediate further 3′-trimming [50].

As mentioned, piRNA pathways differ extremely between species and show a lack of conservation of involved genes [38]. In *Drosophila*, piRNA pathway genes evolve rapidly indicating an arms race between transposons and their cellular defence [5, 37]. In addition, also downstream mechanisms differ between species as *Caenorhabditis elegans* shows an absence of ping-ping amplification using RDR-dependent siRNAs for amplification of the initial piRNAs [16]. Moreover, a screen of non-model species revealed the absence of the piRNA system in nematode lineages other than the most prominent *C. elegans*: these organisms apparently use RDR dependent siRNAs to account for transposon control [46].

Consequently, piRNA biogenesis pathways are highly diverse and increasing evidence indicates piRNAs not restricted to the germline but present in low abundance also in somatic tissues e.g. piRNA-like sRNAs have been identified in various somatic tissues by their ping-pong signature and increasing reports also show piRNAs regulating endogenous genes expression in somatic cells, too [41, 42].

As a result, an ongoing discussion asks for the evolution of these multiple functions of piRNAs and one possibility is the co-option of transposon derived piRNAs to regulate genomic functions [45]. A piRNA analysis of several arthropod species revealed somatic piRNAs targeting transposons and mRNAs across all species and the authors consequently scrutinize that the ancestral role of piRNAs was to protect the germline from transposons [31].

In the context of changing dogmas about Piwis, ciliates provide an excellent model for that as they do not harbor any Agos but contain 15 distinct Piwi proteins [6]. These unicellular eukaryotes undergo sexual recombination of meiotic nuclei in order to develop somatic macronuclei in the same cell with germline micronuclei. Indeed, some Ptiwis show germline specific expression and they were shown to be involved in the accumulation of two classes of meiotic sRNAs involved in elimination of transposons and transposon-derived sequences (IESs) *Paramecium* [12]. Here, we characterize two Ptiwi proteins (Ptiwi13 and Ptiwi14) expressed during vegetative growth. Both have been implicated to be involved in transgene-induced silencing, a mechanism in which introduction of non-expressible transgenes silences an endogenous remote locus [6, 43]. It remains unknown, why two distinct Ptiwi proteins are involved in this mechanism: their involvement and also the 2′-O-methylation of transgene triggered sRNAs [32] is reminiscent of a classical ping-pong mechanism. However this is conflict with the fact that Dicer is involved in the mechanism [15, 30]. One hypothesis may be that both load different classes of siRNAs, because the silenced remote locus was shown to produce secondary siRNAs (2°) as a response to the primary (1°) siRNAs from the transgene [15]. Transgene-induced silencing in *Paramecium* acts on the chromatin level and 2° siRNA appear not to result from full length mRNA rather than nascent transcripts. The aim of this study is to clarify the role and the mechanism of the two distinct Ptiwis in the mechanism of transgene-induced silencing in *Paramecium*.

## Materials and Methods

### 1.1 Cell culture, RNAi, Microinjection

*Paramecium tetraurelia* cells (stock 51 and d4-2) were cultured as described before using *Klebsiella planticola* for regular food in WGP (Wheat grass powder) [48]. All cultures for this study were grown at 31°C. RNAi by feeding of dsRNA producing bacteria was carried out as described before [13, 47] using the double T7 vector L4440 in the RNAse III deficient *E*.*coli* HT115DE3. Microinjection of the pTI-/- and FLAG fusion transgenes was carried out as described before [39].

### 1.2 Phylogenetic analysis

Analysis of Ptiwi amino acid sequence was carried out using MEGAX [28] using the neighbor joining method and 1000 bootstraps replicates. Evolutionary distances were computed using the *poisson* correction method. After pairwise deletion, 962 positions were used for the final dataset. Ptiwi1-15 sequences were described in [6] and we also used a human curated amino acid sequence for the putative pseudogene Ptiwi04 using its paralog Ptiwi05 as a template. Ptiwi16 and and 17 are newly identified Ptiwi proteins (see Suppl. Fig. 1 for sequence, position, and accession number and Suppl. Fig. 2 for a clustalW alignment of all Ptiwi proteins).

**Figure 1.**
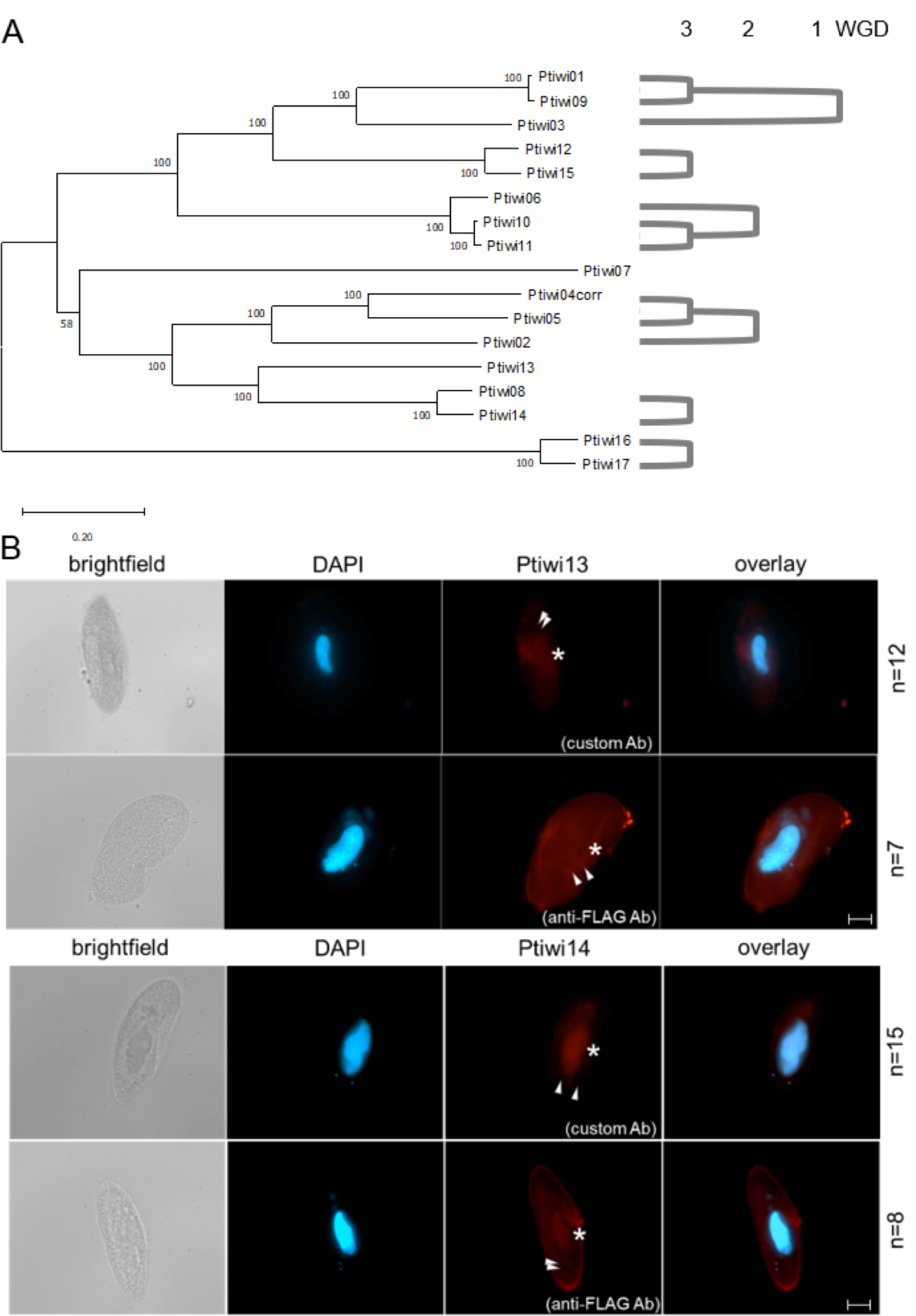
Relationship of Ptiwi proteins and their localisation in vegetative cells. A) Evolutionary relationship of *Paramecium tetraurelia* Ptiwi proteins (strain d4-2) represented by a neighbour joining tree involving 1000 bootstraps replicates (see Methods for details). The amino acid sequence of the putative pseudogene Ptiwi04 was corrected using its paralog Ptiwi05. On the right, ohnologs from Paramecium whole genome duplications (WGD 1-3) are indicated. B) Localisation of Ptiwi proteins in vegetative Paramecium cells injected with Ptiwi13-FLAG (top) or Ptiwi14-FLAG (bottom). Cells were analysed by indirect immunofluorescent staining using custom antibodies directed against Ptiwi13 or Ptiwi14 labeled with secondary Alexa594 - conjugated antibody (red). Additionally, cells were stained using anti-FLAG antibody. Representative overlays of Z-stacks of magnified views are presented. Other panels show DAPI (in blue), brightfield and overlay of DAPI and Alexa594 signal. White arrows point at micronuclei while asterisk is indicating position of the macronucleus. Scale bar is 10 *µ*m and exposure is 2 s. n is number of documented Z-stacks.

**Figure 2.**
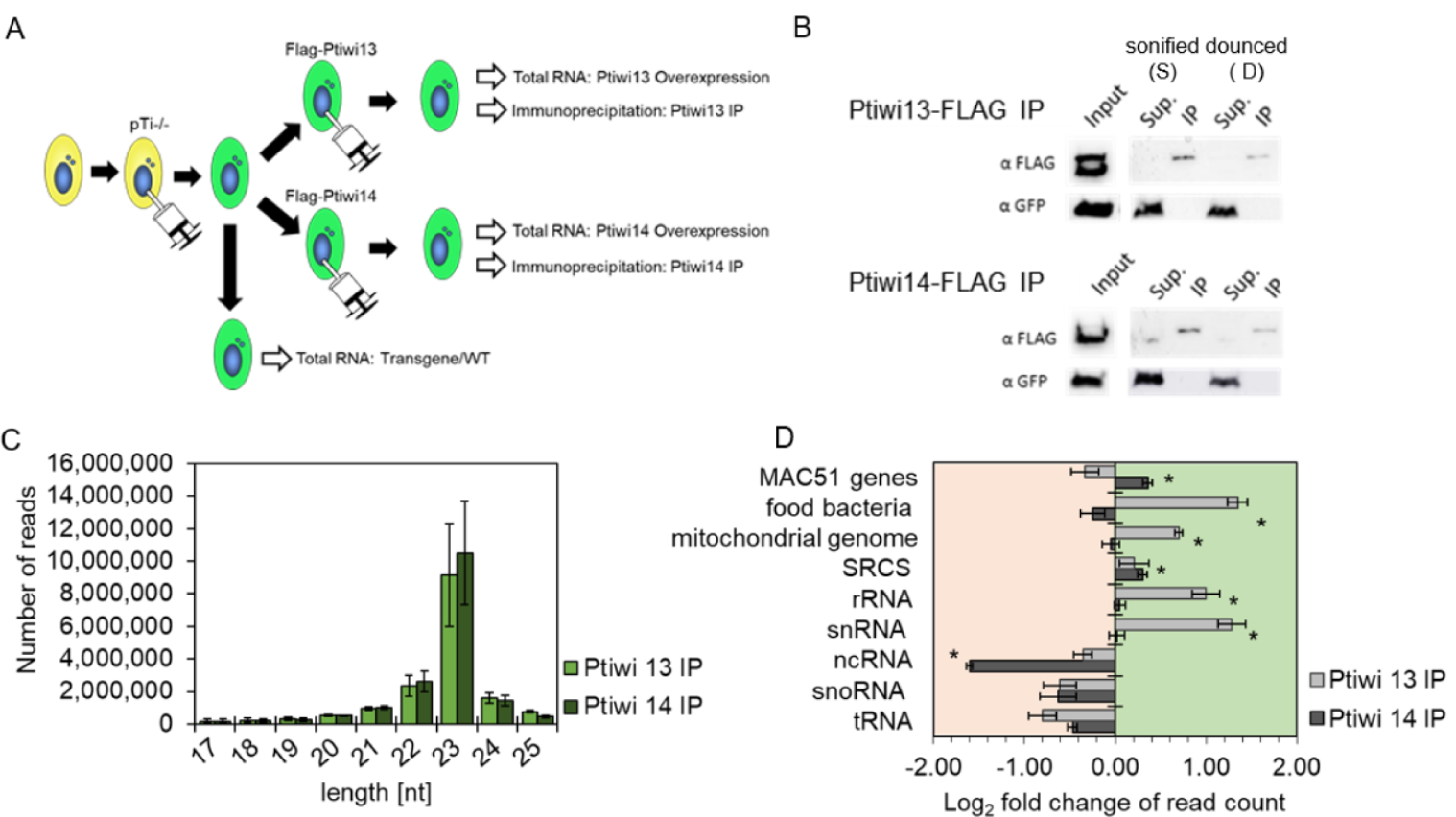
Analysis of sRNAs in Ptiwi immunoprecipitations. A) Experiment overview. A single cell was injected with the pTI-/- transgene and then with FLAG-Ptiwi13/14 constructs, respectively (green). B) Control Western blots for the IPs using anti-FLAG Abs for Ptiwi detection and anti-GFP (Sup.-Supernatant, IP-Immunoprecipitation). Two different setups of the IPs used sonication (S) and douncing (D) for cell lysis, the latter remains MAC structure but permeabilised. C) Total read length distribution of Ptiwi IPed reads after adapter trimming. Reads from three IP replicates each were merged. D) Relative enrichment of RNA reads in Ptiwi IPs mapping to different categories of genomic templates. Enrichment was calculated in reference to individual *PTIWI* overexpressing lines. Reads of three replicates each were merged and compared. * p-value *<*0.005.

### 1.3 RNA isolation and treatment

Total RNA was isolated with TriReagent (Sigma) and integrity was checked by denaturating gel electrophoresis after DNase I (Invitrogen) digestion and subsequent purification with acid phenol. For dissection of 3’-modifications by periodate oxidation, 20*µ*g RNA were dissolved in 17.5*µ*l 4.375mM borax, 50mM boric acid, pH8.6 and 2.5*µ*l 200mM sodium periodate were added. After 10min incubation in the dark, 2*µ*l glycerol were added with another 10min incubation. After drying in the speedvac, the pellet was dissolved in 50*µ*l 33.75mM Borax; 50mM boric-acid; pH 9.5 and incubated for 90min at 45°C. The RNA was subsequently purified with Sephadex G-25 columns (GE).

### 1.4 sRNA sequencing and analyses

For siRNA sequencing, 17-25 nt small RNA fractions were isolated by denaturating PAGE and subjected to standard small RNA library preparation using the NEB Next small RNA sequencing Kit (NEB, Frankfurt a.M., Germany). The procedure includes 3′-OH and 5′-monophosphate specific ligation steps and we tried to lower 3′-2′-O-me biases by 18 hours 3′-ligation at 16°C. After 10 PCR cycles, the libraries were gel-purified and sequenced on the HiSeq 2500 using the Rapid Mode with 28 cycles. Reads were de-multiplexed and adapter sequences were trimmed using Trim Galore (http://www.bioinformatics.babraham.ac.uk/projects/trimgalore/) that uses Cutadapt [33] with a stringency cutoff of 10. For analysis of reads, we used normalised counts, and converted these values to Transcripts Per Million (TPM), which we also refer to as sRNA accumulation. For the analyses specific to endogenous clusters, a TPM value greater than one was termed to be present in Ptiwi IPs. We used the RAPID pipeline to obtain the normalised counts, implementing the KnockDown Corrected Scaling (KDCS) method [24]. Hierarchical clustering of data sets was performed with complete linkage using an euclidean distance measure and heatmaps were created using R/Bioconductor package gplots (v3.0.1.1). Data is deposited at the European Nucleotide archive, ENA, Acc. Nr. PRJEB38766.

### 1.5 sRNA signatures

Sequence logos of 23nt sRNAs were generated using WebLogo3 [10] with error bars twice the height of the correction for small sample size. Probabilities for overlapping reads from aligned sRNA reads were calculated using the small RNA signature analysis tool in Galaxy [2]. sRNAs from 17 to 25 nt were mapped to each region of interest, allowing no mismatches or multimapper in bowtie [29] and overlaps from 1 to 25 nucleotides were calculated. Plots for read length distribution and coverage were created using Geneious Prime 2020.1.2.

### 1.6 Antibodies, Western Blots, Immunostaining

Peptides corresponding to the amino acids 684-698 and 449-463 of *Paramecium* Ptiwi13 (C-DDAPPQARKNNKSPY) and Ptiwi14 (C-QNWMQRLTAEIGDK) respectively were used for immunisation of rabbits. Purification of antibodies from serum was performed by coupling the respective peptides to SulfoLink*™* coupling resin (Thermo Scientific) and following the manuals instructions. Purified antibodies were tested by dotblot assays (see Suppl. Fig. 3). Western Blots were carried out as described previously [27] using indicated antibodies diluted 1:250 in 5% milk/TBST. Indirect immunofluorescence staining was carried out as previously described [11]. Cells were permeabilised in 2.5% Triton X-100 and 1% formaldehyde for 30min followed by fixation in 4% formaldehyde and 1.2% Triton X-100 for 10min. After blocking in 3%BSA/TBST cells were incubated in primary antibody diluted 1:200 in 3%BSA/TBST under mild agitation over night at 4°C. After washing and incubation with 1:2500 Alexa Fluor ® 568 F(ab’)2 fragment of goat anti-Rabbit IgG (H+L) (Thermo Scientific # A-21069) cells were stained with DAPI and mounted in VECTASHIELD (VectorLaboratories). Images ware acquired using Zeiss Axio Observer with ApoTome. For expression of tagged Ptiwis, the respective orf was cloned into *Paramecium* FLAG-Vectors pPXV containing three FLAG sequences either at the N-or C-terminus (kind gift of M. Valentine and J. Van Houten, Vermont, USA) as described in [53]. Injected clones were screened by single cell PCR and positives were grown for cell fixation and protein isolation. Macronuclei were isolated as described [40] and protein was isolated by adding preheated Laemmli sample buffer with subsequent boiling for 5 min.

**Figure 3.**
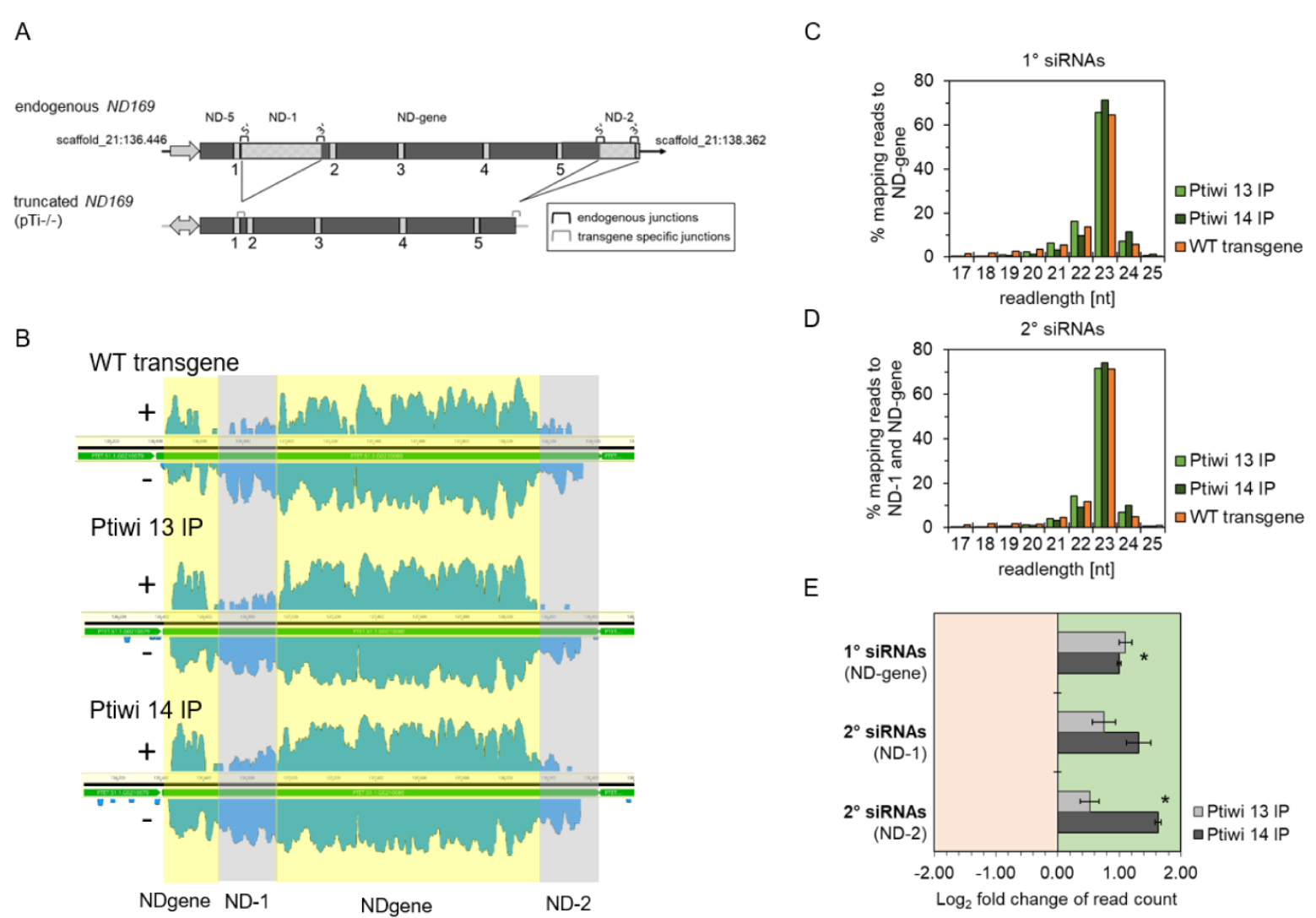
Ptiwi13 and 14 load transgene-associated sRNAs. A) Detailed scheme of the endogenous *ND169* locus (top) and the truncated transgene (bottom). Introns are numbered and brackets symbolise specific junctions. Shaded regions are not part of the transgene (ND-1 and ND-2). B) Coverage tracks of siRNAs mapped to the endogenous ND169 locus. siRNAs were separted by their direction (sense/antisense). Regions accounted for 1° siRNAs (NDgene, grey) and 2° siRNAs (ND-1 and ND-2, yellow) were highlighted. Coverage in log scale is shown for one replicate each. C) Read length distribution of 1° siRNAs and 2° siRNAs from Ptiwi IPs and control (WT transgene). Data is shown as proportion of reads mapping to the NDgene locus and D) the ND-1 and ND-2 locus. E) Relative enrichment of RNA reads in Ptiwi IPs mapping to different regions of the transgene. Enrichment was calculated for merged reads of three replicates in reference to individual Ptiwi overexpressing lines. * p-value *<*0.005.

### 1.7 Ptiwi-Immunoprecipitation

Transgenic Ptiwi lines harboring the pTI-/- transgene and a single Ptiwi-FLAG fusion construct (described above) were used for Ptiwi IPs using monoclonal anti-FLAG M2 (Sigma). Our procedure follows the protocol by [12] for developmental Ptiwis with the following modifications. 500k cells of a single transgenic line were grown and harvested by snap freezing in 2ml lysis buffer. 1ml of the lysate was broken in a Dounce homogeniser and 1ml was sonified until also Macs were destroyed. After addition of 50*µ*l Alexa Fluor ® M2 Magnetic Beads (Sigma) and incubated over night by gentle agitation. After washing beads with wash buffer and re-suspended in 100*µ*l. 10*µ*l were used for western controls by addition of 2.5*µ*l Laemmli sample buffer and subsequent boiling for 2min. 90*µ*l were extracted with TriReagent LS (Sigma) according to the manufacturers recommendation.

## 2 RESULTS

## 2.1 Ptiwi13 and 14 are polyphyletic slicers in cytoplasm and nucleus

In this study, we focus on the relationship between Ptiwi13 and Ptiwi14 which have been implicated to be involved in transgene-induced silencing [6, 15]. It remains unclear why two different Ptiwis are involved in this pathway. We aimed to answer the question whether ping-pong amplification contributes to sRNA accumulation or whether e.g. these two Ptiwis have different loading preferences for 1° and 2° siRNAs.

Figure 1A shows the evolutionary relationship between the 17 identified Ptiwi proteins. The analysis indicates the transgene related Ptiwis 13 and 14 not to be related and chromosome synteny analysis [3] does not suggest them to be the result of a whole genome duplications of which the *Paramecium tetraurelia* genome underwent at least three [4]. Thus, both appear to be polyphyletic. In addition, Suppl. Fig. 2 suggests that both Ptiwis are capable of Slicer activity due to the presence of the catalytic DDH triad in the C-terminal area. We raised antibodies against specific peptides corresponding to both Ptiwis for immunolocalisation (Suppl. Fig. 3 and Methods). Fig. 1B indicates broad cytosolic Ptiwi13 signals and a clear absence in the macronucleus while staining of Ptiwi14 shows weak enrichment in the macronucleus. Please be aware that this cannot demonstrate absence of the respective protein in any compartment. The sub-cellular localisation are supported by amino-acid analysis by the ngLoc method (Suppl. Fig. 4) [26]. We conclude that both Ptiwis have different sub-cellular localisation preferences.

**Figure 4.**
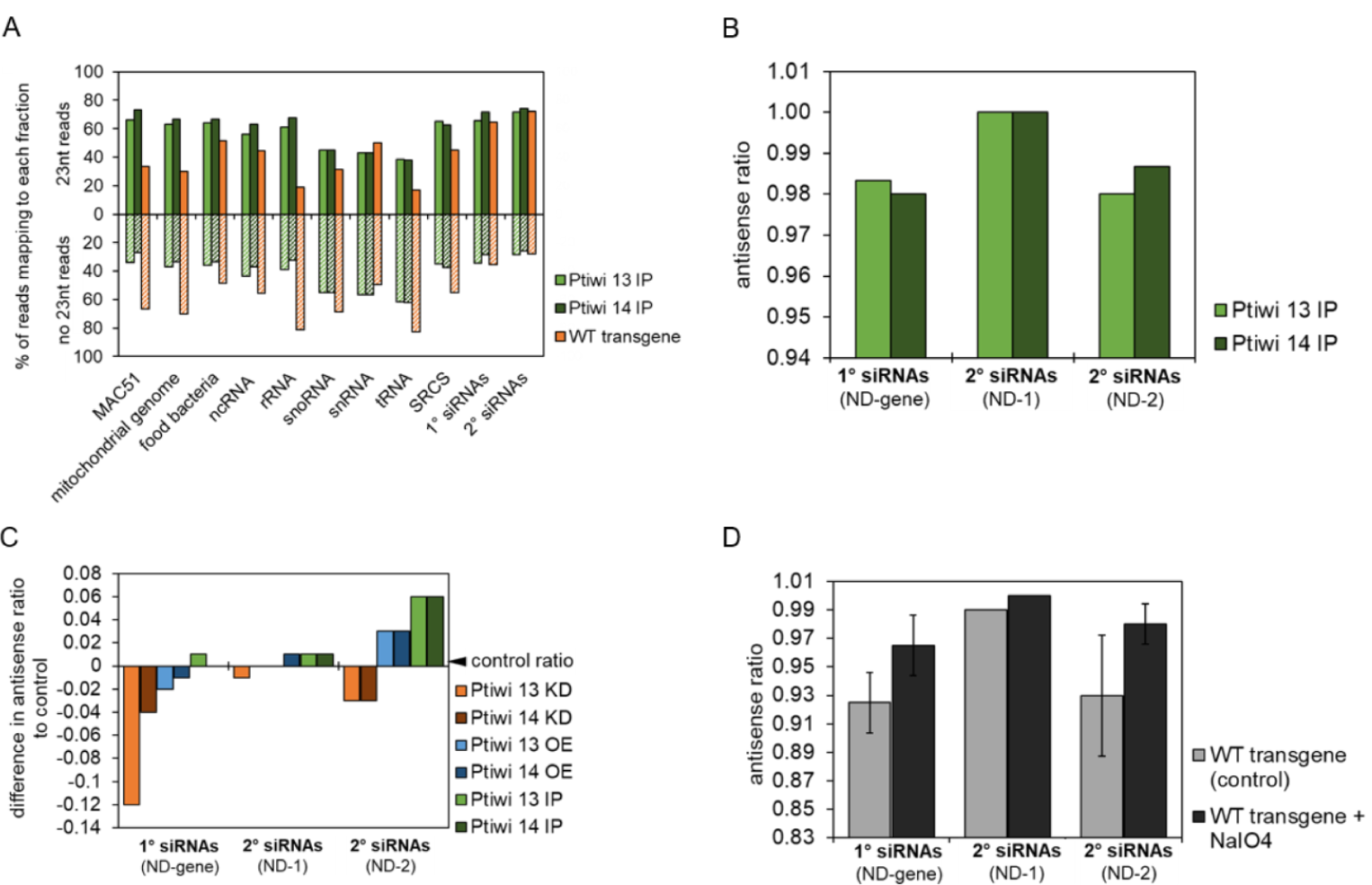
Asymmetric modification and Ptiwi selectivity contributes to accumulation of antisense siRNAs. A) Number of 23nt reads maping to each indicated genomic feature and transgene regions accounting for 1° and 2° siRNAs were related to the total number of reads of other sizes. Calculation is shown for the mean of Ptiwi IPs and WT transgene (control). B) Antisense ratio of reads from Ptiwi IPs calculated by merging three replicates each. C) Difference in the antisense ratio of reads to the antisense ratio of respective control was calculated. Data is shown for reads from knockdown of Ptiwis, overexpression and IPs (S) mapping to the indicated transgene regions. D) Antisense ratio of small RNAs from WT transgene samples (untreated) and small RNAs treated with sodium periodate (+NaIO4). Reads of two replicates were merged.

## 2.2 Ptiwi13 and 14 have different loading preferences for endogenous and exogenous sRNAs

For Ptiwi IPs, we injected Flag-tagged Ptiwi13 and 14 transgenes respectively into a transgenic strain harboring the pTI-/- transgene line (Fig. 2A). This transgene contains GFP marker and additionally a truncated version of the endogenous ND169 gene causing silencing of the endogenous remote locus. Western blots of aliquots of the lysates/pulldowns verified the successful immunoprecipitaion and the absence of soluble proteins present in the supernatant (Fig. 2B). As a first insight, the trimmed read length distribution of Ptiwi IPed reads shown in Fig. 2C reveal a clear 23nt peak which is the predominant siRNA read length in *Paramecium*. We then mapped reads to different classes of RNA templates and quantified them relative to the respective abundance in *PTIWI* overexpression lines. Fig. 2D indicates that Ptiwi13 enriches for sRNAs of exogenous precursors such as food bacteria and mitochondria. This is in agreement with a previous reports that Ptiwi13 is also involved in endogenously triggered RNA by bacterial dsRNA and that *Paramecium* also converts exogenous ssRNA into siRNAs [6, 8]. In addition, Ptiwi13 also enriches for fragments of rRNA and snRNA. In contrast, Ptiwi14 IPs show accumulation of transgene-associated sRNAs and other siRNA producing genes (SRCs) of the *Paramecium* genome [23]. Our data indicates that Ptiwi14 exclusively loads these two classes and interestingly both are also enriched in Ptiwi13. The data shows that both Ptiwis are clearly not redundant but have different localisation and loading preferences.

We further dissect the transgene associated sRNAs in the Ptiwi IPs to get an insight which sRNAs are specifically loaded into these Ptiwis.

### 2.3 2° siRNAs are enriched but not exclusively found in Ptiwi14

Both Ptiwis have been earlier shown to be necessary for efficient transgene-induced silencing [6]. Fig. 3A shows the genomic structure of the endogenous *ND169* gene involved in trichocyst discharge. This gene becomes silenced on the chromatin level when cells are injected with a truncated form of this gene shown below: the pTI-/- transgene shows two deletions: one on the 5’-coding region (ND-1) and the 3’-coding region including the 3′-UTR (ND-2) [15]. Mapping sRNA reads to the endogenous *ND169* as shown in Fig. 3B therefore shows regions specific for 2° siRNAs as these are deleted from the transgene: siRNAs mapping to these regions, ND-1 and ND-2, result the endogenous *ND169* gene, thus being per definition 2° siRNAs [15]. Those regions existing in the transgene and the endogenous gene (called NDgene in the following) consist of both, 1° and 2° siRNA, however as the abundance of 2° siRNAs is more than 10fold lower compared to 1°, the NDgene regions shows predominantly 1° RNAs. Fig. 3B also shows the reads from Ptiwi IPs. These maps already rebut one of our first hypothesis on the question why two different Ptiwis are involved in this mechanism: the data clearly shows that both Ptiwi IPs show reads mapping to the NDgene and to the ND-1/2 region. As such, they do not mutually exclusively load 1° siRNAs and 2° siRNAs. Analysing quality and quantity of these sRNAs, Figs. 3C and 3D show that both Ptiwi bound classes are of predominant 23nt length. Ptiwi14 significantly enriches more 2° siRNAs (Fig. 3E). The finding that silencing of the *ND169* gene by the transgene was shown to occur on the chromatin level may make sense in this context as 2° siRNAs are then produced from nascent transcripts in the nucleus and loaded by nuclear Ptiwi14.

### 2.4 Ptiwi13 and 14 specifically load 23nt antisense siRNAs

Still, our data does not allow for a conclusion why two Ptiwis are involved in this mechanisms and what the precise role of those could be. To follow this question, we had a look for the ratio of 23nt reads to other read lengths. Comparing bulk RNA to IPs, Fig. 4A shows that both Ptiwis specifically load 23nt sRNAs, however the ratio of 23nt to other lenghts varies between different RNA species. For several RNAs, e.g. food bacteria, rRNA etc. one can identify that Ptiwis specifically select 23nt sRNAs among many other RNAs. It seems likely that fragments of these RNAs are produced by different mechanisms creating several lengts of sRNA of which Ptiwis enrich for 23nt sRNAs.

This appears different for transgene associated 1° and 2° siRNAs which show almost identical ratio of 23nt siRNAs in bulk RNA and IPs, suggesting that a distinct biogenesis mechanisms contributes to more precise sRNA cleavage. Going more into detail with these transgene associated siRNAs, both Ptiwis load predominantly antisense RNAs as shown in Fig. 4B which could be an argument against ping-pong amplification. Comparing the antisense ratio of *PTIWI* knockdown and IPs to each other, our data indicates a certain decrease of the antisense ratio on knockdowns and an increase in IPs (Fig. 4C). These changes in the antisense ratio are only moderate, and it is either possible that both Ptiwis complement for each other to some extent or that also other factors contribute to strand selection. In many systems, 2’-O-methylation was shown to occur in context of Piwi associates sRNAs. Using periodate oxidation of RNA and subsequent library preparation, we can show that both, 1° and 2° are resistant to periodate thus likely to be methylated at the 3’-end (Suppl. Fig.5). Our data moreover indicates that predominantly antisense sRNAs are modified in this manner (Fig. 4D) suggesting that this modification contributes to stand selection and stabilisation.

### 2.5 Ptiwi14 loaded siRNAs have a 5’-Uridine preference

As the current data implicates that both Ptiwis select strand from Dicer cuts rather than amplify sRNAs in a ping-pong manner, we followed this idea by analysing the sequence logos of transgene associated sRNAs. Fig. 5A shows logos of 1° and 2° sRNAs obtaining 5’-Uridine preference in bulk RNAs. To decide whether these RNAs should result from a Dicer cut one should see a A preference at position 21 for the non 5’-U reads. These are shown in Fig. 5A, but one cannot identify such a Dicer signature nor for a ping-pong signature. Ptiwi IPs (Figs. 5B, C) reveal that the 5’-U preference of bulk RNA is mainly due to Ptiwi14 whose RNAs show a much stronger 5’-U preference compared to Ptiwi13. Unfortunately lack of Dicer or ping-pong logos do not allow for further conclusions about the biogenesis mechanisms.

**Figure 5.**
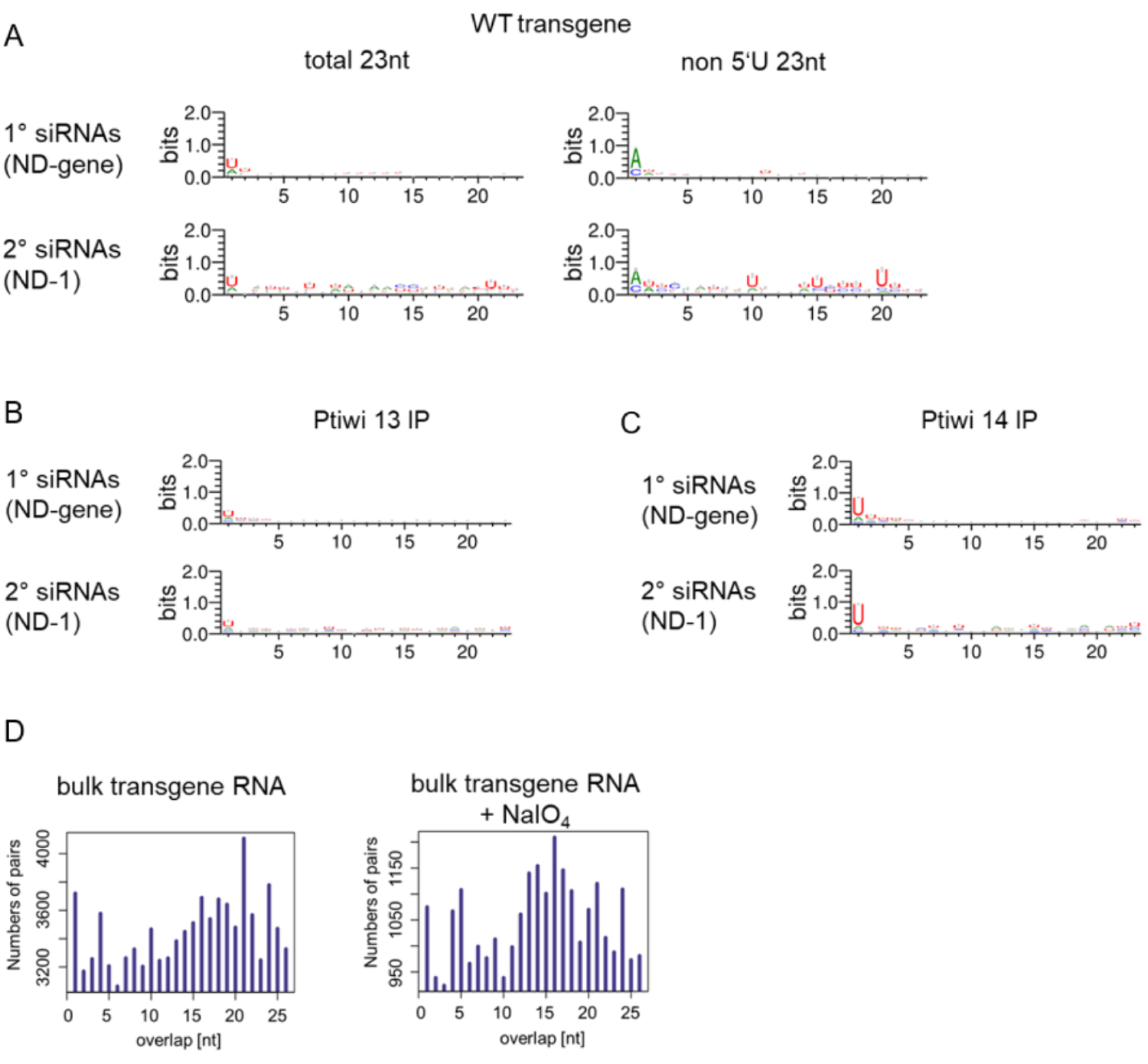
Sequence logos of 1° and 2° siRNAs. A) Sequence logos of 23nt antisense reads mapping to transgene regions. Logos for either all sequences or the ones without 5’-U are shown. B) Sequence logos of 23nt antisense reads from Ptiwi 13 IP (S) and C) and Ptiwi 14 IP (S). D) Overlap predictions of small RNAs from 17 to 25 nt of wildtype transgene RNA (untreated) and the same small RNAs treated with sodium periodate (+NaIO_4_).

We therefore additionally analysed reads for overlapping reads: in this analysis transgene 1° siRNAs show a clear peak at 21nt overlaps which fit to 23nt Dicer cuts (Fig. 5D). Interpreting this as a argument for Dicer cleavage, this is contrary with the missing Dicer signature in sequence logos. We have to consider, that the observed 5’-U preference is not that strong compared to 5’-Us in scnRNAs for instance [30, 44], and thus the complementary 21-As on the passenger strand might also be less significant. In addition, Suppl. Fig. 5 shows that the 5’-U preference is still pronounced in periodate treated RNAs, thus indicating that nucleotide preference and methylation co-occur on the same molecules. Fig. 5D in addition shows that we cannot identify any 21nt read overlaps in periodate treated samples in agreement with the hypothesis of strand specific methylation.

### 2.6 sRNA Uridine content contributes to strand selection

When analysing the sequence logos, not only the 5’-U preference was observed as some logos suggested that stabilised strands are rich of Uridines (Suppl. Fig. 6, 7). Because the logo analysis is not suitable for the analysis of general nucleotide composition, we followed this by analysis of the antisense ration along the transgene and endogenous *ND169*. Fig. 6A shows again that most areas show dominant antisense preference for 1° and 2° siRNAs: an exception is the promoter proximal region (called ND-5) which shows almost 50/50 strand distribution. Dissecting these different regions we consequently calculated the Uridine and Adenosine content of these regions (Fig. 6B) revealing that the ND-5 region is different to the other regions as it shows a much higher Uridine content. We therefore asked whether this could be seen in sRNAs, too. Also in sRNAs the ND-5 regions shows a different behaviour compared to other regions (Fig. 6C) and we consequently calculated the U-content of 23nt sRNAs of (i) *in silico* diced RNA (ii) bulk transgene siRNAs, and (iii) siRNAs of Ptiwi IPs. As demonstrated in Fig. 6D, the exceptional sense ration of the promoter proximal ND-5 regions is due to the enrichment of U-rich sRNAs, mainly by Ptiwi14. The analysis further reveals first that for all regions, the U-content of the more abundant antisense RNAs is higher than the sense siRNAs and second for almost all regions for 1° and 2°, the U-content of Ptiwi IPed sRNAs is above *in silico* diced sRNAs. Thus, strand selection by Ptiwi13 and Ptiwi14 seems to include a general selection for Uridine rich sRNAs.

**Figure 6.**
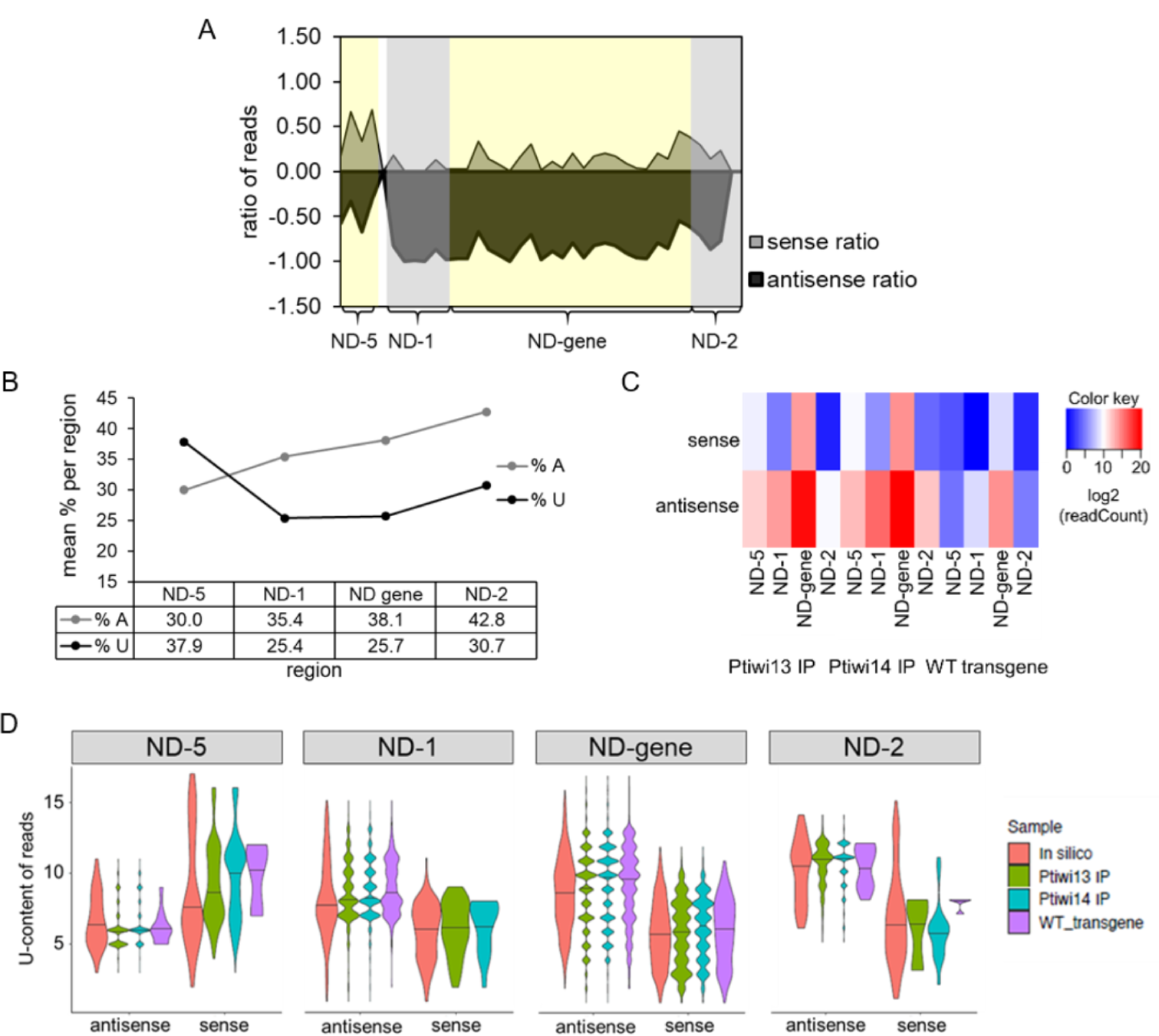
U-content analysis of 23nt sRNAs in Ptiwi IPs. A) Sense/antisense reads of different regions in 50nt windows. B) Percentage of A and U nucleotides of the sense RNA transcript of each region. C) Heatmap of reads mapping to the indicated regions separated by their direction. D) X-axis shows U-content of reads found in Ptiwi IPs while density represents number of reads. In silico data is generated by counting U-content of 23mers of the DNA sequence for each region.

**Figure 7.**
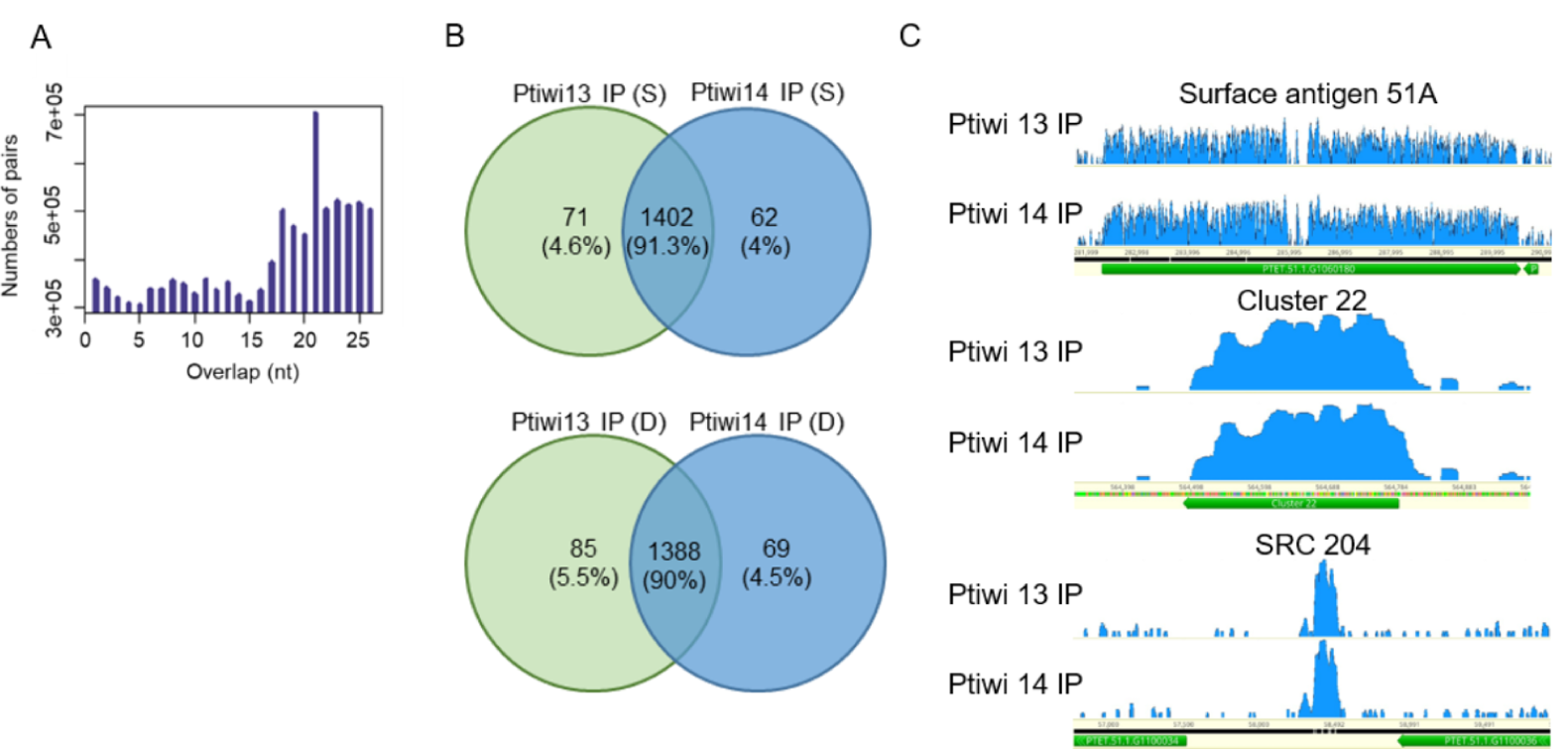
Endogenous sRNAs in Ptiwi IPs. A) Calculation of overlaps of endogenous 17-25nt sRNAs isolated from wildtype RNA. B) sRNAs mapping to SRCs (small RNA clusters) in the *Paramecium* genome are analysed by their presence in Ptiwi IPs. Venn diagrams of the two IPs of sonified (S) and dounced (D) lysates with numbers of SRCs detected and the proportion of each fraction of the total found SRCs. C) Three examples of endogenous loci shown by coverage tracks of unnormalised data (surface antigen 51A, cluster22 and SRC 204).

### 2.7 Transgene-induced silencing mimics endogenous siRNA accumulation

We finally had a look for endogenous siRNAs. We have recently described siRNA producing loci in the *Paramecium* genome showing read length preference of 23nt [23]. Figure 7A shows also for these endogenous clusters a predominant overlap of 21nt indicating Dicer to be involved at least in the majority of them. More surprisingly most of the endogenous clusters can be identified also in both IPs of Ptiwi13 and 14 (Fig. 7B/C and Suppl. Fig. 8). As a result, transgene-associated silencing appears to have common genetic requirements with endogenous siRNA accumulation pathways and appears therefore as a suitable model to study siRNA accumulation.

**Figure 8.**
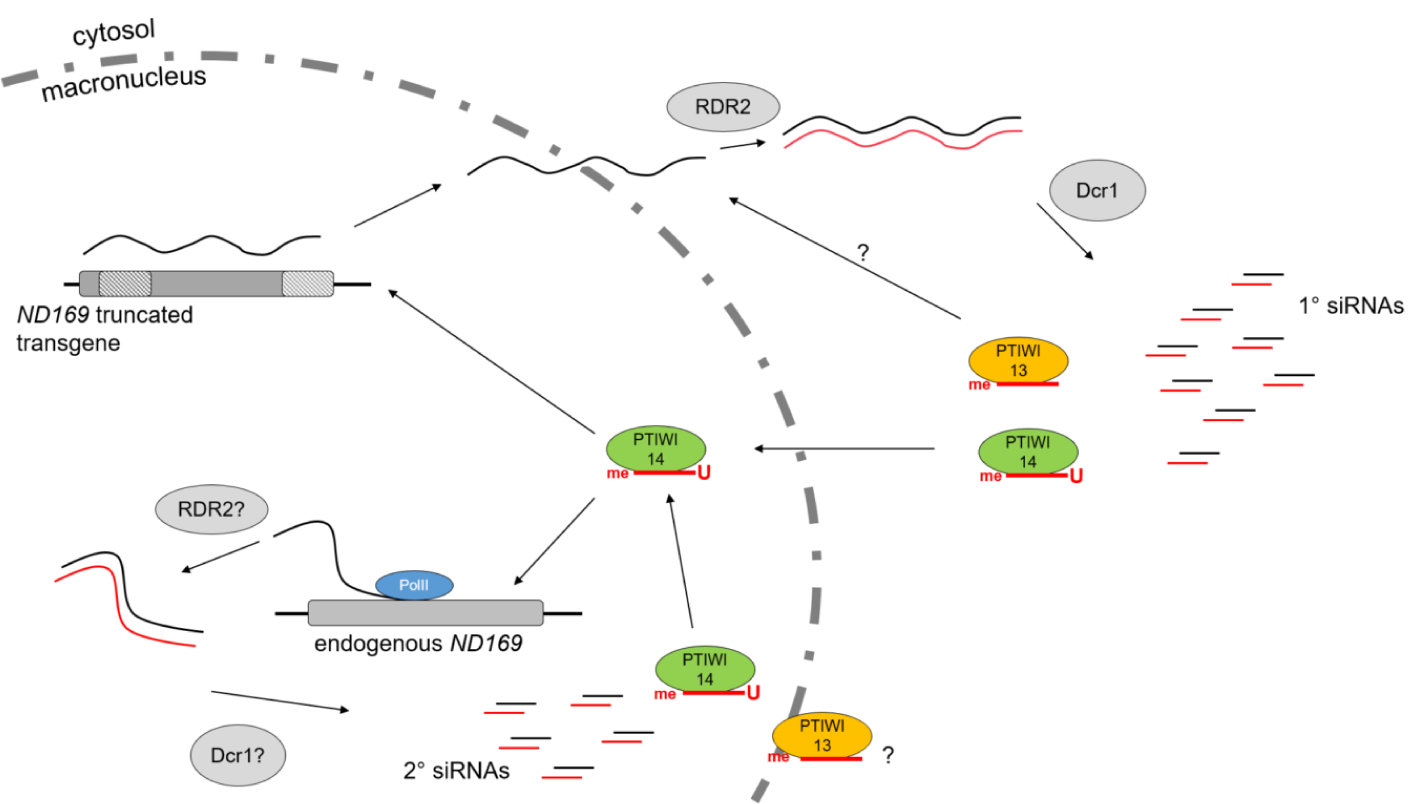
A working model for transgene-induced silencing in *Paramecium* based on this and earlier studies. RDR2 and Dicer1 produce 1° siRNAs from truncated transgene transcripts and Ptiwi13/14 select antisense strands which show 3’-methylation. Transport into the nucleus occurs via unknown shuttling mechanisms. Targeting nascent transcripts at the remote locus, chromatin modification (not shown) and 2° siRNAs are triggered. 2° siRNAs are likely produced by similar mechanisms as 1° siRNAs and antisense strands are loaded into Ptiwi14.

## 3 DISCUSSION

The dissection between Agos and Ptiwis was originally not only based on sequence homology but their spatial and temporal activity in germline and somatic cells. The most important distinction between both sRNA loading complexes was the aspect due to their action in strand selection: Agos load RNAse III generated duplexes whereas Piwis load longer ssRNA and generate their own sRNAs. Although many recent reports were published describing non-canonical functions of Piwis in somatic cells target non-transposable elements, the latter aspect of strand selection activity remains an important difference between the two groups. Ciliates belong to the chromalveolates which are equidistant to animals, plants and fungi [25] and especially *Paramecium* is different to many other species, not only metazoans, as the genome does not contain Agos, but 15 Ptiwis [6] and our work adds two more (Ptiwi16 and 17). Thus, *Paramecium* offers exciting possibilities for evolutionary comparison of RNAi mechanisms and the mode of action of individual components.

### 3.1 Transgene associated siRNAs are Dicer products

We started this work here based on the surprising finding that two different Ptiwis are involved in a process where a non-expressible transgene silences a remote locus at the chromatin level [6, 15]. The first logical idea why two Ptiwis would be involved in this mechanism would be the action of ping-pong amplification. Although this would have been supported as Ptiwi13 and Ptiwi14 own Slicer activity, it is in conflict with the fact, that Dicer1 is necessary to produce at least the 1° siRNAs. The second logical hypothesis would have been, that both Ptiwis distinguish between 1° and 2° siRNAs. Our data clearly shows that both hypotheses are not true, because we do not see any ping-ping signature in either 1° and 2° siRNAs but a 21nt overlap of reads thus strongly suggesting that Dicer cuts at least the majority of siRNAs rather than Ptiwi slicing. Although each individual knockdown of *PTIWIS* shows effects on 1° and 2° siRNAs thus not allowing for a clear conclusion about their specification, Ptiwi IPs disprove also the second hypothesis, because both Ptiwis load both, 1° and 2° siRNAs, however in different quantities.

But what is the function of both Ptiwis then? Could they have the very same function being redundant? This seems not very likely because our data shows some discrete differences between Ptiwi13 and Ptiwi14: (i) Ptiwi13 loads more sRNAs from exogenous templates, (ii) both have different sub-cellular localisation, (iii) Ptiwi14 loads more 2° siRNAs and finally (iv) Ptiwi14 shows a much stronger preference for 5’-U RNAs. It seems therefore more likely that both Ptiwis have indeed distinct and specialised functions in this mechanism.

Concerning the different subcellular localisation, we need to interpret fluorescence data with care, as we cannot exclude that also fewer portions of Ptiwis are in other cellular compartments. It seems more likely that both are involved in a shuttling process between cytosol and nucleus, reminiscent of the nematode Ago NRDE-3 which localises in the cytosol and redistributes to the nucleus when bound to 2° siRNAs from the feeding pathway [17]. This would make sense if we think about the trigger for transgene-induced silencing. It has been shown, that explicitly non-expressible transgenes induce RNAi [13, 14] tempting that a quality control mechanism is involved in this process to dissect which transgene can produce translatable mRNA. These processes, e.g. nonsense mediated RNA decay, work in the cytosol and as we have previously shown that transgene-induced silencing works on the chromatin level, this cytosolic signal needs to be transported into the nucleus. Right now we cannot reveal the precise function of both Ptiwis but a role in a transport process seems more likely than stationary Ptiwis. First of all, because both Ptiwis contain 2° siRNAs: these show decreasing coverage along the endogenous target genes [15] and previous data indicates them to be produced from unspliced precursor RNA: this kind of a nascent transcript will likely stay in the nucleus and therefore, we hypothesise that 2° are produced in the macronucleus (Fig. 8). Our data does not allow for a conclusion why we find 2° siRNAs in cytosolic Ptiwi13.

### 3.2 Ptiwis select strands from Dicer products

Our data indicates that both Ptiwis select for strand specific sRNAs from Dicer cut duplexes which represents a non-canonical function of Piwi proteins. This finding is fostered by several aspects. First, Dicer knockdown reduces all sRNAs [15, 30] and in addition we have shown here bulk transgene siRNAs show 21nt overlaps of 23nt RNAs. Our study also brings strand asymmetry in association with individual properties of sRNAs apparently contributing to strand selection and stabilisation by Ptiwis: 5’-U preferences, U-content and 3’-methylation. 5’-U preferences have been frequently described, e.g. for the Piwi lacking *Arabidopsis* Ago1 [35]. It is also quite reminiscent to the strong 5’-U preference of *Drosophila* Piwi (and weaker in Aubergine) which act together with Ago3 in the ping-pong amplification of piRNAs [7]. However, we cannot identify any sRNAs with an A-preference at position 10 which would result from such a mechanism.

5’-nucleotide preferences were also reported also for the developmental Ptiwis in *Paramecium* showing a strong 5’-UNG signature [12, 30, 44]. However, recent evidence from *in vitro* dicing experiments show that this signature is due to cleavage preference of the involved Dicer-like enzymes rather than due to preferential Ptiwi loading [18]. In this context, the authors speculate about a co-evolution of Dicer-like enzymes to produce sRNAs targeting germline specific DNA with strong sequence bias at its ends. This seems likely for the particular need to target conserved sequence-ends, but for the control of endogenous gene expression accumulation of such conserved sequence features in siRNAs would not make sense, especially as our data indicates this mechanisms to be quite similar to the endogenous siRNA pathways.

Ptiwi IPs show that both Ptiwis select for 23nt siRNAs which is the predominant length of somatic sRNAs in Paramecium [23, 30], but this is not the case for all Ptiwis as developmental Ptiwi09 does not show a clear length preference for sRNAs [12]. In transgene-induced silencing, it seems likely that the 5’-U preference is due to Ptiwi selection rather than a Dicer1 preference because the preference differs between both Ptiwis.

General preferences for nucleotides apart of the ultimate 5’-end have also been reported for human miRNAs, where purines (A/G) are enriched in the guide strand likely interacting with aromatic residues in the PAZ domain of Agos and the passenger is therefore rich in pyrimidines (U/C) [20]. Other preference have been reported for plant Ago2 and Ago5 which show strong bias for either adenine or cytosine [51]. Our observed U-preferences of the two *Paramecium* Ptiwis complements the knowledge of general nucleotide preferences in contrast to sequence signatures and may be useful knowledge to design more powerful silencing constructs for *Paramecium*.

However, it is also clear now that nucleotides as a biochemical reason and also the resulting thermodynamic behaviour of a sRNA duplex are only individual aspects for strand selection. It has been demonstrated that many protein factors additionally contribute to strand selection in addition to Agos allowing for dynamic adaptation of the miRNA system in response to challenges to adapt gene expression [34]. Another aspect which needs to be taken into account is the availability of RISC targets. It has first been shown in plants that the alteration of the target RNA binding quality alters miRNA abundance [52]. Having the target effect in mind, this can also explain the strand selection in association with 3’-methylation in association with Ptiwi13 and 14: our results show that the methylation occurs predominantly on antisense strands which let us hypothesise, that the selection of strands is carried out by Ptiwis, but for the later methylation, factors of available target RNAs may play also a role. This modification has until now not been described for any other sRNA in *Paramecium* but is well known for piRNAs where the 3’-end is methylated after trimming. As we can rule out this mechanism, Ptiwis apparently select for the guide strands of Dicer cleaved duplexes and our data let us hypothesise that they interact with the Hen1 methyltransferase for strand specific methylation by the fact that antisense ratio in periodate treated samples is always higher than in untreated samples. This is a clear difference to plant Hen1 activity on siRNAs duplexes [21] and would be more similar to animal Hen1 activity on ssRNA stimulated by Ago/Piwi interaction.

## 4 CONCLUSION

Our data here suggests a mechanism involving Dicer cleavage and subsequent strand selection by two distinct Piwi proteins. This is unexpected, because strand selection from double stranded siRNA duplexes is usually done by Agos, whereas Piwis use their slicer activity for target RNA cleavage and ping-pong amplification of sRNAs. In addition, most Piwis have been described to be meiosis specific rather than somatic. Both Ptiwis here work apparently in an Ago mode and select 23nt guide strands due to 5’-U and total U-content (Fig. 8). This finding is another evidence for the extreme diversification of small RNA amplification not only in ciliates. In particular, our data in line with reports of other species about many non-canonical functions of Piwi proteins, which let us scrutinise whether a canonical function of those can be defined. Agos are absent in ciliates and the broad diversity of Piwi function has already been demonstrated by the differential usage of two sets of Ptiwis for scnRNAs and iesRNAs which can be seen as two steps of 1° and 2° developmental sRNAs involved in elimination of germline DNA. In this mechanism, 1° and 2° developmental sRNAs have been shown to be loaded in distinct Ptiwis, contrary to our vegetative mechanism which does not involve a preferential loading of 1° and 2° transgene-induced siRNAs into Ptiwi13 and 14. The knockdown of Ptiwi10 and Ptiwi09 during meiosis resulted in accumulation of duplexes and the authors concluded that these Piwis could be responsible for strand selection in reminiscence of Agos. This is further supported by the involvement of Dicer/Dicerl-like proteins in the biogenesis of these two sRNA classes [44] thus indicating that the two developmental Piwi proteins act also more like Agos than ping-pong related Piwis which is similar to our finding here. Thus, *Paramecium* appears to use Piwi proteins indeed more in a Ago-manner, as shown for the totally different mechanisms of germline IES excision and vegetative chromatin silencing. Especially the comparison between different ciliates, e.g. *Tetrahymena* and *Paramecium*, reveals unexpected diversity of Piwi related siRNA mechanisms as for instance secondary (2°) sRNAs from excised genetic regions become produced before excision in *Tetrahymena* and after excision from circular concatamers in *Paramecium* [1, 36]. Both mechanisms in addition do not have any features of a ping-pong mechanism, thus changing our point of view on the variety of Piwi functions. Ciliates may in general use Piwis in an Ago manner which is surprising not only for the vegetative Ptiwis described here but even more for the massive elimination of transposons and transposon-derived sequences during development which is the hall mark of Dicer independent piRNA in other organisms. Our data raises then the question whether ciliates use their Piwis totally different to other species or whether this function of PIWIs could be the primordial one and later split into Agos and Piwi mode of actions. However the latter would be in conflict with the finding that even bacteria have Ago proteins [22], although their function is less understood.

## Acknowledgments

This work was supported by grants from the German research Council (DFG) to MS (SI1379/3-1), MHS SCHU (3140/1-1) and MJu MJ (CRC894).We are grateful to Dominique Furrer and Mariusz Nowacki, Bern, for sharing RIP experience, Kaz Mochizuki, Montpellier, for advice in Ptiwi-IPs, and Megan Valentine and Judy VanHouten, Vermont, for sharing the FLAG fusion vector.

