## Supplementary Data for "Two Piwis with Ago-like functions silence somatic genes at the chromatin level"

### Suppl. Fig 1.

#### Ptiwi16 is GSPATG00013703001

>Ptiwi16

MSGTIVVGSNIYETEFLDEIFIHYGNKTADTISFKDGLIISPKKLELPEYKINRSEQGEV

IMAILKEAMNLLEYSKLSDLGIINMFELQYVNQLKSYPSLRFQFMDKEQPSILMDVRKSY

FTVHNCYQIIKTLNTEPSAYFKNKFVLLRYPEKKMIVQIKSVVVDQEFTYKGVSLEDYFK

QKYQLKSKSSVYLETYNTQNIKKQFLIAEFCDLKIDYIEQDATVYCQQLEYWNKIINNSS

SFIKFLNQYKLIIRNEQVEFKQQYLEPGNLVLNNQTKSIFQIVASRTSSEFEYYYTDFFL

KNASTFTTSLTINKLIILVTREITELKDNFNIFLEKFDLFIKSRQYKISQPEIYDINNNL

NEILEKYVTEDSYVLFVGDQKTDFTSARNTLLSKAIPNQLINLPIQDQEINRLLAIMTAN

LGSVPWSIKEINGQINNKKSAVLGIWKSDNSFSACLSINKYLNKYISSFGDLDQILTLLL

STYYGTHKTLANQIIIFAEEDYTDIMEQRIQGLIQLIQDQGLAESTQKPEIVMVQVKDAS

EERYFTYFERNFESYYCKNPQVGCIVPINEKQGRFSFHSQRGEQGCLQAQQLQISSSNRR

EIAQLYYHLIFLNFDLSSITYLPAPLHYAKKLSKHQLQSEQYKKAIEKGHMLFV

#### Ptiwi17 is the second part of GSPATP00016985001 as this automatically annotated genes was artificially fused with the 5´-upstream ORF

#### The corrected amino acid sequence is:

>Ptiwi17

MSGTIVVGSNIYETEFLDEIFIYYGNKTNNTIGFKDGLIISPKKMELPEYKMNRSERGEI

IMAIIKEAMNLMEYSKLSDMGIINMFELQYINQLKSYPSLRFQLMDREQPSVLMDVRKSY

FTMHNCYQIIKTLNTDPSTYFKNKFVLLRYPEKKMIVQIKSVVLDQEFTYKGLSLEDYFK

QKYQLRSKSSVYLETYNTQNIKKQFLIAEFCELKIDYIEQDVTVYCQQLEYWNKIINNSS

SFIQFLNQYKFIIRNEQVEFKQQYLEPGNLALNNQTKSIFQIVASRTSSEFEYYYTDFFL

KNPTSFTTSLTINKLMILVTREIIELRDNFKSFLEKFDLFIKSRQYKISQPQIYDINNNL

NEILEKYVTEDAYVLFVGDQNTDFIQARNTLLSKAIPNQIIILPIQDQEINRLLAMMTAN

LGSVPWAIREINGLITNKKSAVLGIWKSENSYSACLSINKYLNKYISSFGDLDQVLTLLL

ATYYATHKILANQIIVFAQDDYTQIMEQRILGVIQLIRDQGLAQQILKPEVVMVQVKDAT

AERYFTYFERNFESYYCKNPQVGCIIPINEKQGRFSFHSQRGEQGCLQAQQLHISSTNTK

EIAQLYYHLIFLNFDLSSTTYLPAPLHYAKKLSKHQIQSEQYKKAIEKGFLLFV

### Suppl. Fig 2.

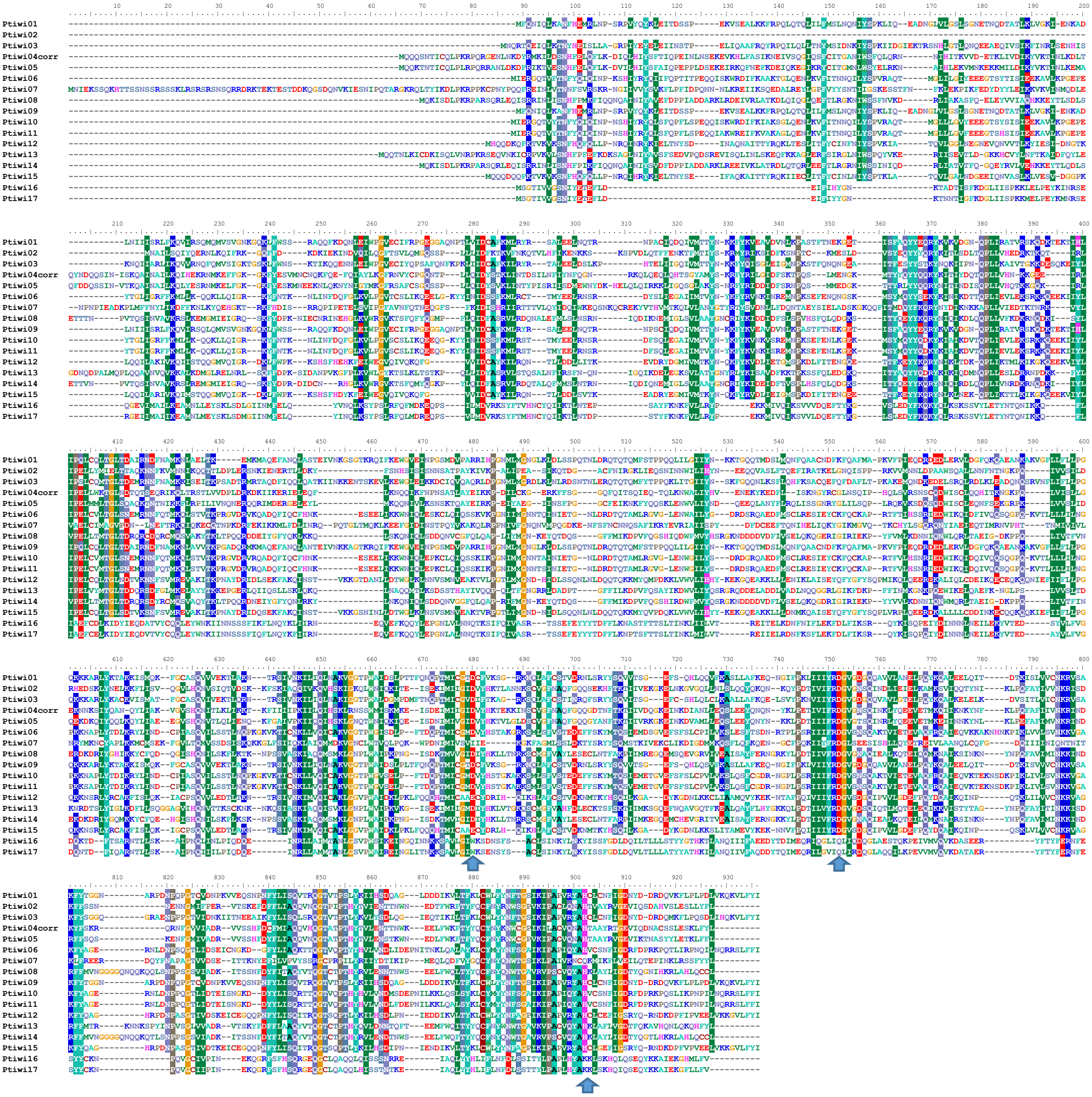

**Suppl. Fig. 2** ClustalX alignment of the 17 Ptiwi Amino Acid Sequences of *P. tetraurelia*. For Ptiwi04 the gene sequence was artificially corrected to a translatable orf using the paralog Ptiwi05 as a template. Blue arrows indicate conserved catalytic residues of the DDH triad.

Suppl. Fig. 3

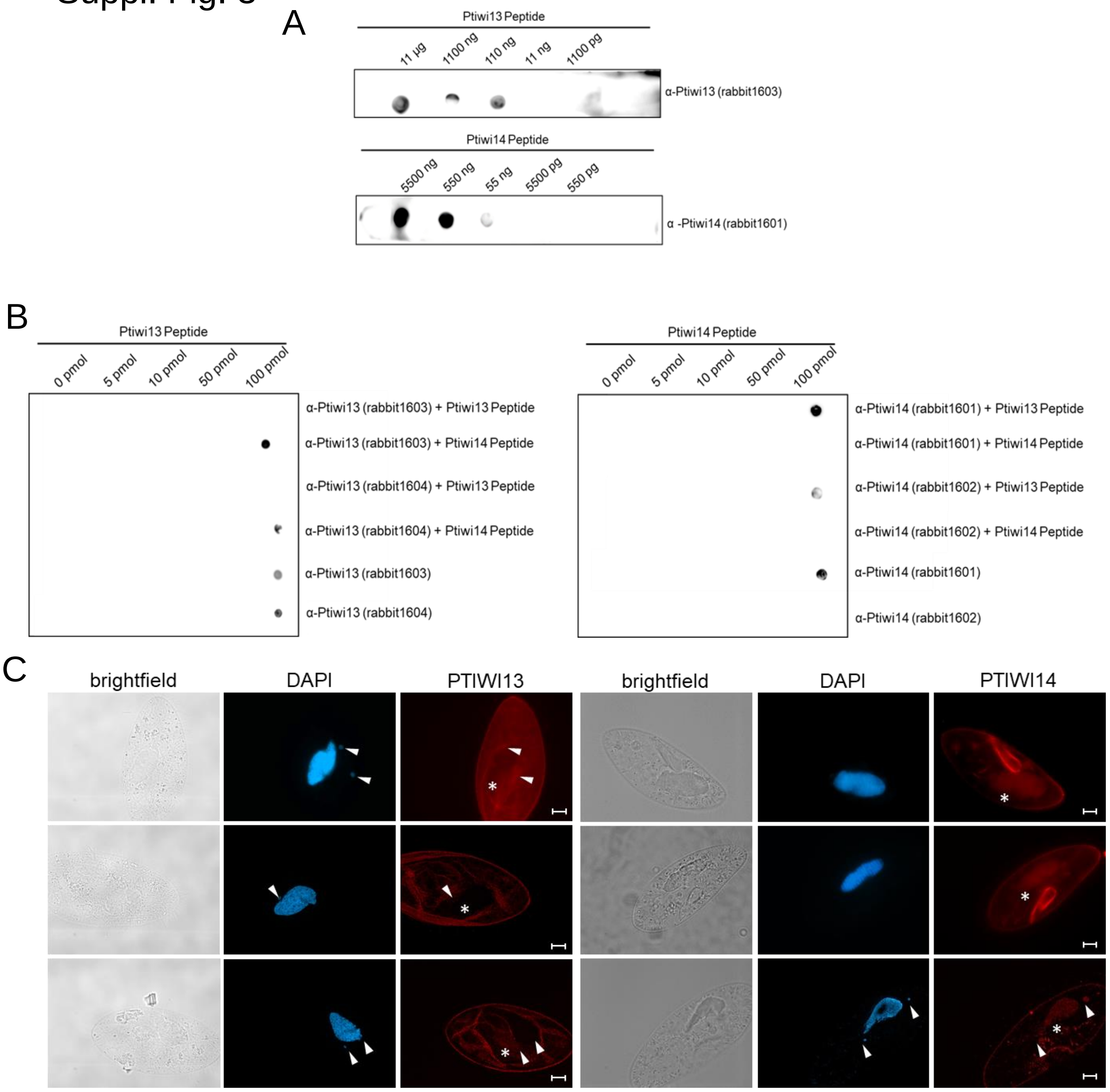


**Suppl. Fig. 3** Validation of antibody specificity. A) Dot blot assay using custom made antibody directed against *P. tetraurelia* Ptiwi13 and Ptiwi14. Different amounts of each indicated peptide were spotted on the membrane. Hybridization with the respective antibodies showed the reactivity with the corresponding peptide. B) Competition assay. 0 – 100 pmol of each peptide were spotted. The antibodies were mixed with the corresponding peptide in 100x excess in advance for 1 hour, then the mix was used to decorate the membrane. Blocking α-Ptiwi13 with Ptiwi13 peptide results in loss of antibody binding on the membrane, while blocking with the Ptiwi14 peptide does not have an effect on the signal (mock). Same is true for validation of Ptiwi14 antibody (bottom right).C) Localisation of Ptiwi proteins in vegetative Paramecium cells. Cells were analyzed by indirect immunofluorescent staining using custom antibodies directed against Ptiwi13 and Ptiwi14 labeled with secondary Alexa594 - conjugated antibody (red). Representative overlays of Z-stacks of magnified views are presented. Other panels show DAPI (in blue) and brightfield. White arrows point at micronuclei while asterisk is indicating position of the macronucleus. Scale bar is 10 μm and exposure is 2 s.

Suppl. Fig. 4

| ID | Prediction | Pred1 | Prob1 | Pred2 | Pred2 | Pred3 | Prob3 | MLCS |
|---|---|---|---|---|---|---|---|---|
| PTWI14 | NUC | NUC | 10.45 | CYT | 10.32 | CSK | 9.444 | 20.74 |
| PTIWI13 | CYT | CYT | 13.67 | NUC | 11.35 | CSK | 9.118 | 24.44 |

NUC - Nucleus
CYT - Cytoplasm
CSK - Cytoskeleton
MLCS - multi-localization confidence score

**Suppl. Fig. 4** Prediction of protein sub-cellular localisation of Ptiwi13 and Ptiwi14 using ngLoc method.

Suppl. Fig. 5

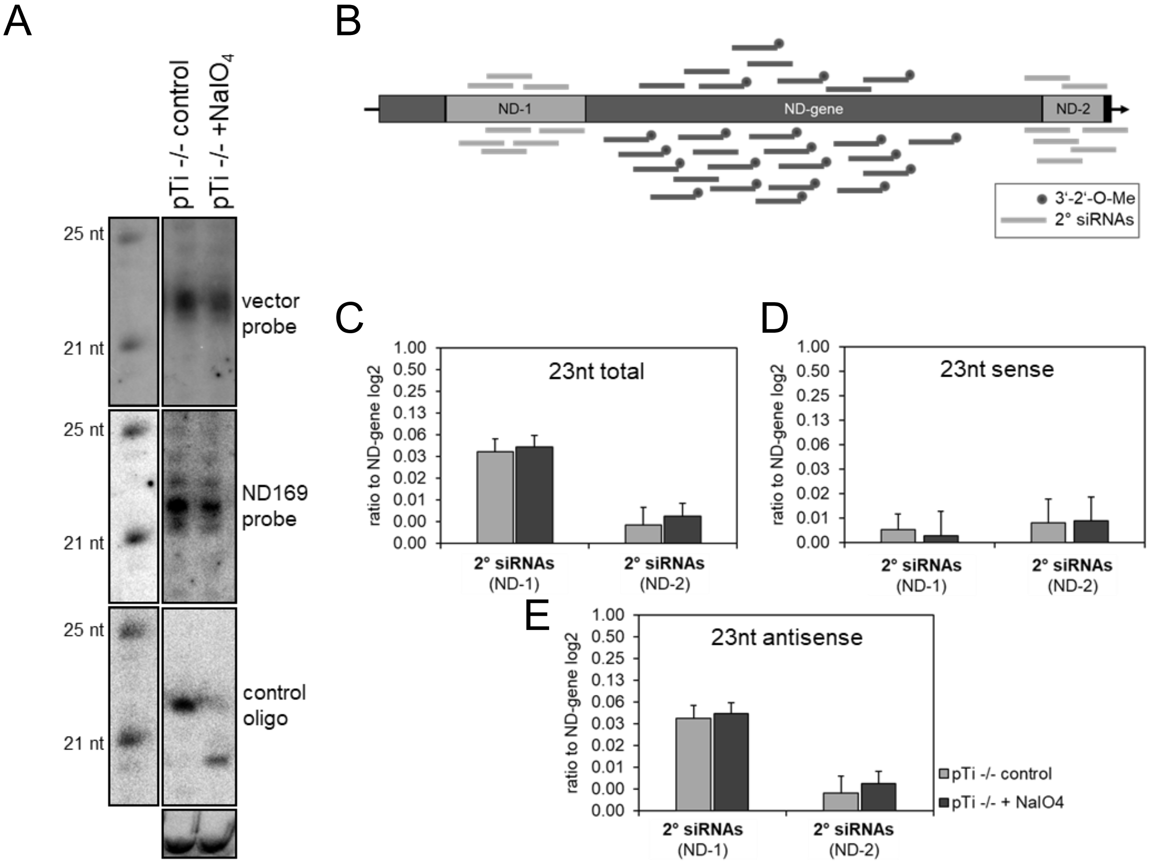


**Suppl. Fig 5** Analysis of 3´- modifications of siRNAs. A) Northern blot analysis of transgene siRNAs. Untreated RNA and $NaIO_4$ treated RNA (pTI -/- + $NaIO_4$) was blotted. Next to the marker on the left, the blot was hybridized with a transgene vector probe (outside *ND169* region, upper blot), a ND169 probe (middle) and a as probe against the 3´- unmodified spike in oligo (lower blot). B) Simplified scheme of *ND169* regions and normalization procedure visualized by arrows. Light grey regions are not part of the transgene and account for production of secondary sRNAs. C-E) Figs. show results of sRNA seq of untreated and $NaIO_4$ treated RNA normalized to the total number of 1°-*ND169* mapping siRNA. Data of two replicates were merged.

### Suppl. Fig. 6

A

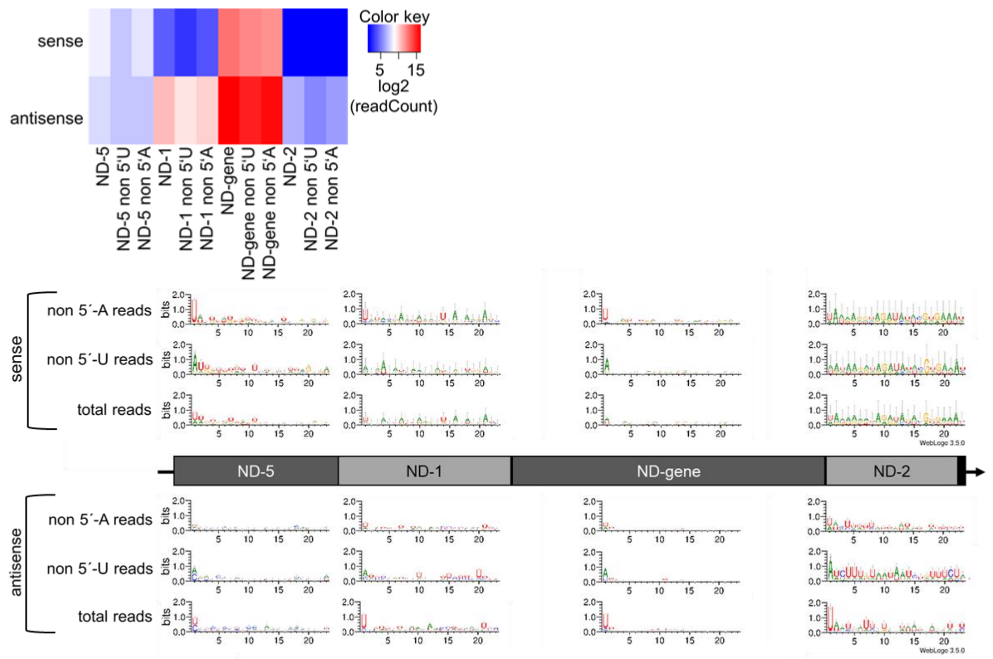


B

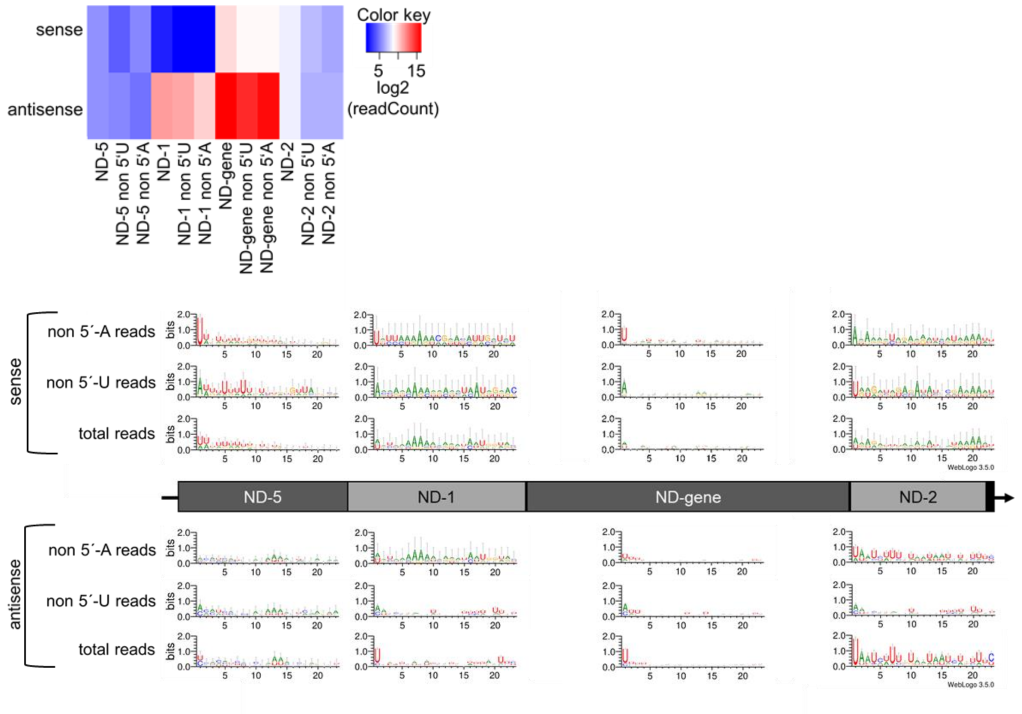


**Suppl. Fig. 6** Sequence logos 1° and 2° siRNAs**.** Sequence logos of 23nt siRNAs in A) ICL KD and B) ICL KD periodate treated RNA are shown. Heat maps show log2 of number of reads from each sample used for plotting the different logos.

Suppl. Fig. 7

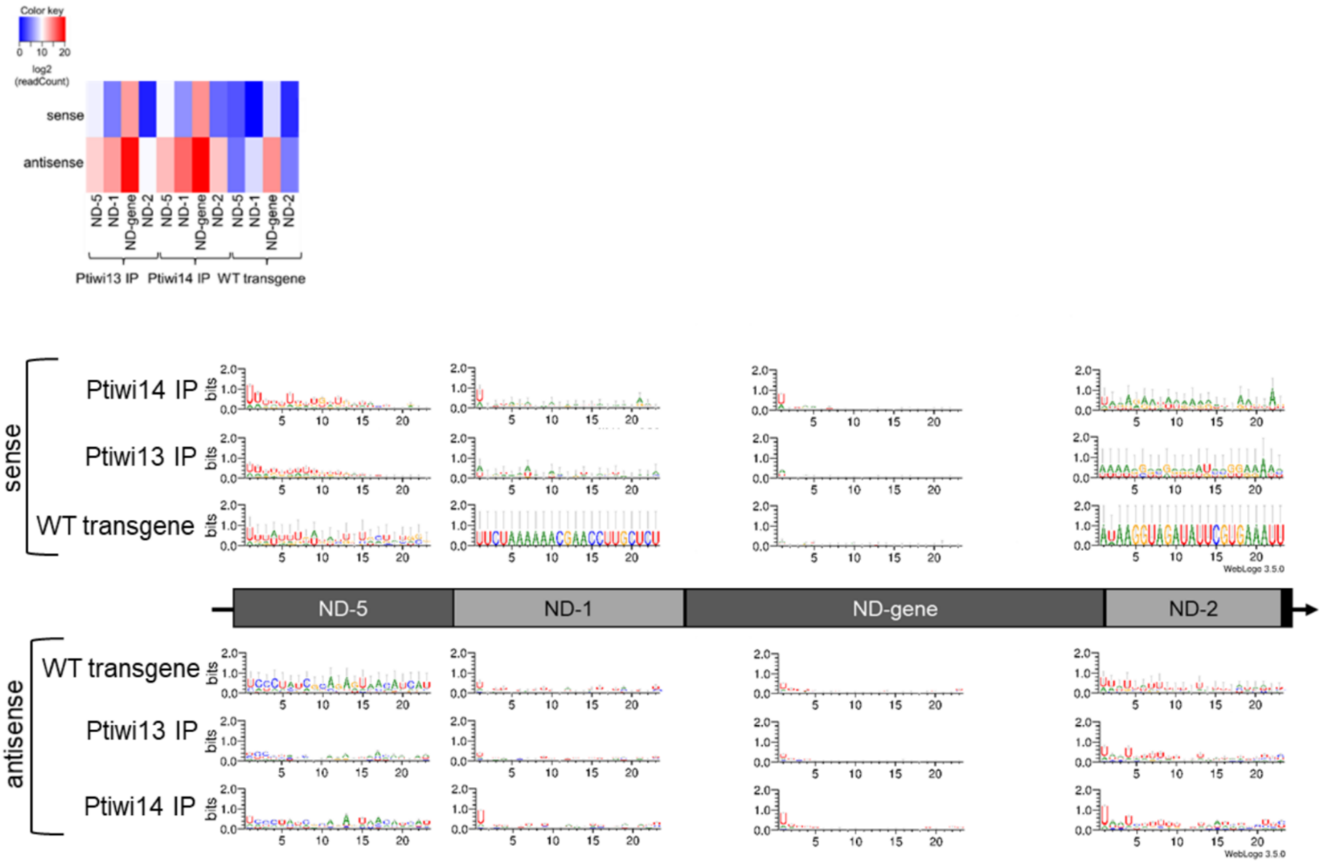


**Suppl. Fig. 7** Sequence logos of 23nt siRNAs in Ptiwi IPs. Sequences were separated by their direction (sense, top; antisense, bottom). Heat maps show log2 of number of reads from each sample used for plotting the different logos.

Suppl. Fig 8.

A B

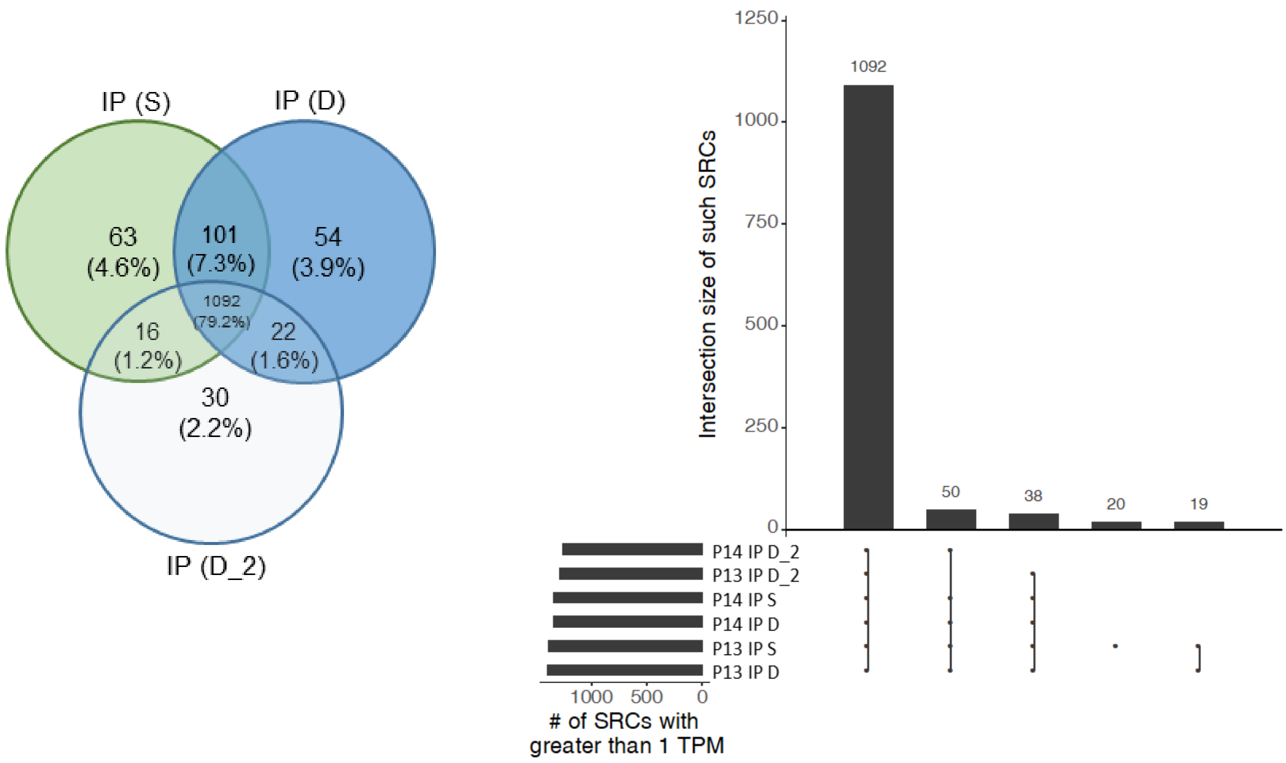


**Suppl. Fig. 8**. Analysis of Ptiwi IPs for presence of endogenous sRNAs. Samples used for this comparison are S (sonified lysate), P1 (dounced lysate with intact but permeabilized Macs) and P2 (dounced lysate with subsequent sonficiation for Mac destruction). A) shows a Venn diagram indicating the number of SRCs (endogenous small RNA clusters) appearing in ≥1TPM in IP libraries. Below the percentage of covered SRCs is indicated. B) Set intersection plot of the same data.
